# Actomyosin-driven force patterning controls endocytosis at the immune synapse

**DOI:** 10.1101/452896

**Authors:** Anita Kumari, Pablo J. Sáez, Mathieu Maurin, Danielle Lankar, Mabel San Roman, Raphael Voituriez, Katharina Hennig, Vanessa F. Boura, Mikael C.I. Karlsson, Martial Balland, Ana-Maria Lennon Dumenil, Paolo Pierobon

## Abstract

An important channel of cell-to-cell communication is direct contact. The immune synapse is a paradigmatic example of such type of interaction: it forms upon engagement of antigen receptors in lymphocytes by antigen-presenting cells and allows the local exchange of molecules [1]. Although [2], how forces organize and mechanics has been shown to play an important role in this process impact on synapse function is unknown. We found that mechanical forces are spatio-temporally patterned at the immune synapse: global contractile forces are observed at the synapse periphery and local point-like forces are detected at its centre. The global contractile forces result from a pulsatile centripetal actomyosin flow that leads to formation of F-actin protrusions from which the central point like forces emerge. Noticeably, these force-producing actin protrusions constitute the main site of antigen extraction and endocytosis. Accordingly, deletion of the myosin IIA gene leads to impaired B cell responses. The interplay between global and local forces governed by the actomyosin cytoskeleton therefore controls the endocytic function of the immune synapse and might constitute a more general mechanism in the physical regulation of cell-cell interactions.

---

Cells are endowed with the ability to internalize substrate-bound molecules, which they recognize through specific surface receptors. While the role of substrate mechanics has been extensively investigated in the context of adhesion, its impact on receptor endocytosis remains unclear. A typical case of coupling between substrate mechanics and juxtacrine signalling (i.e. by direct contact) occurs at the immunological synapse, i.e. the tight contact zone that forms between a lymphocyte and an antigen-presenting cell [1], [3]. In the case of B lymphocytes, synapse formation results from the engagement of the B-cell antigen receptor (BCR) by antigens exposed at the surface of neighbouring cells *in vivo*. The immune synapse provides a platform that facilitates signalling and leads to antigen internalization within B cells [4]-[6] which is needed for B cells to ultimately produce high-affinity antibodies and generate immune memory ([7], [8]). As, in general, endocytosis often involves surface-tethered rather than soluble molecules when occurring in tissues, antigen internalization at the B cell synapse provides a valuable model to study the impact of mechanics in this process.

Different experimental systems have been developed as surrogate antigen-presenting cells to study antigen extraction at B cell synapse *ex vivo*: planar lipid bilayers [9], plasma membrane sheets [10] and polystyrene beads [11]. On bilayers, the immune synapse consists of a set of concentric patterns in which molecules and cytoskeletal components are partitioned: a distal supramolecular antigen cluster (dSMAC) with an actin ring, a peripheral supramolecular antigen cluster (pSMAC) enriched for adhesion molecules and a central supramolecular antigen cluster (cSMAC) in which antigens concentrate [9]. The first antigen extraction model to be proposed was based on the observation that B cells spread over antigen-coated substrates and then contract [12], allowing the transport of BCR-bound antigens towards the cSMAC. A second model arose from Atomic Force Microscopy experiments monitoring interactions between the BCR and plasma membrane sheet-bound antigens: they showed that B cells actively pull on BCR-antigen complexes, pulling being required for antigen internalization [10]. Both these mechanical models rely on the same actin based molecular motor, non-muscular myosin II, which in one case is responsible for global cell contraction and antigen transport towards the cSMAC through the generation of actin flows [13], [14], whereas in the other case it was proposed to directly pull on individual BCR-antigen complexes for internalization. However, it is difficult to compare the two models as (1) they involve surrogate substrates with different physical properties (lipid bilayer versus plasma membrane sheets) and (2) they are based on experiments performed at different spatial scales (cell versus molecules) and temporal scales (minutes versus seconds).

Here we investigated the spatio-temporal organization of forces exerted by B lymphocytes during antigen extraction. We show that they display a stereotypical patterning profile that includes two components: (1) peripheral forces resulting from the centripetal flow of myosin II and (2) central forces exerted by local actin protrusions. Strikingly, we found that these actin protrusions need myosin II-dependent peripheral forces to form and are responsible for antigen extraction, reconciling the two models that had been previously proposed. We conclude that the interplay between global and local forces governed by the dynamics of the actomyosin cytoskeleton controls endocytosis at the immune synapse. Myosin II-dependent force patterning therefore emerges as a key event in the regulation of cell-cell interactions.

We used time-dependent traction force microscopy (TFM, see Mat. and Meth.) [15]—[18] to analyse the spatio-temporal distribution of forces at the B-cell synapse **(Fig. 1a)**. Primary naive B cells freshly purified from the spleen of mice expressing a hen egg lysozyme (HEL)-specific BCR were plated on gels coated with HEL or with bovine serum albumin (BSA) as a negative control. A rigidity of 500 Pa was selected for the gels, as this matches the physiological rigidity of macrophages that present the antigen to B cells *in vivo* [19]. Surprisingly, scanning electron microscopy (SEM) analysis showed no B cell spreading on antigen-coated gels, spreading being observed on glasses coated with equivalent amounts of antigen, as previously reported **(Fig. 1b)**. Instead, when B cells contacted the antigen-coated gels, they exhibited pulsatile contractions (**Movie 1**). To characterize this cell mechanical behaviour, we quantified the stress **(Fig. 1c)** and the strain energy exerted by B cells on the substrate. We found that the strain energy displayed a growth phase lasting ~5 minutes, followed by a plateau **(Fig. 1d,e)**. The growth phase was barely observed in the absence of HEL **(Fig. 1d, e, Supp. Fig. 1B, Movie 2**) and the plateau displayed a clear dose dependence on this antigen **(Fig. 1f)**. Measurement of the bead displacement field flux through the cell boundaries showed that the forces detected were mostly directed inward **(Fig. 1g)**, indicating that they corresponded to pulling rather than pushing forces. Consistent with this result, small gel pieces appeared to be pulled by the protrusions that formed at the lymphocyte periphery **(Fig. 1b)**. These results show that B cells exert contractile pulling forces on substrates of physiological rigidity in an antigen-dependent manner.

**Figure 1:**
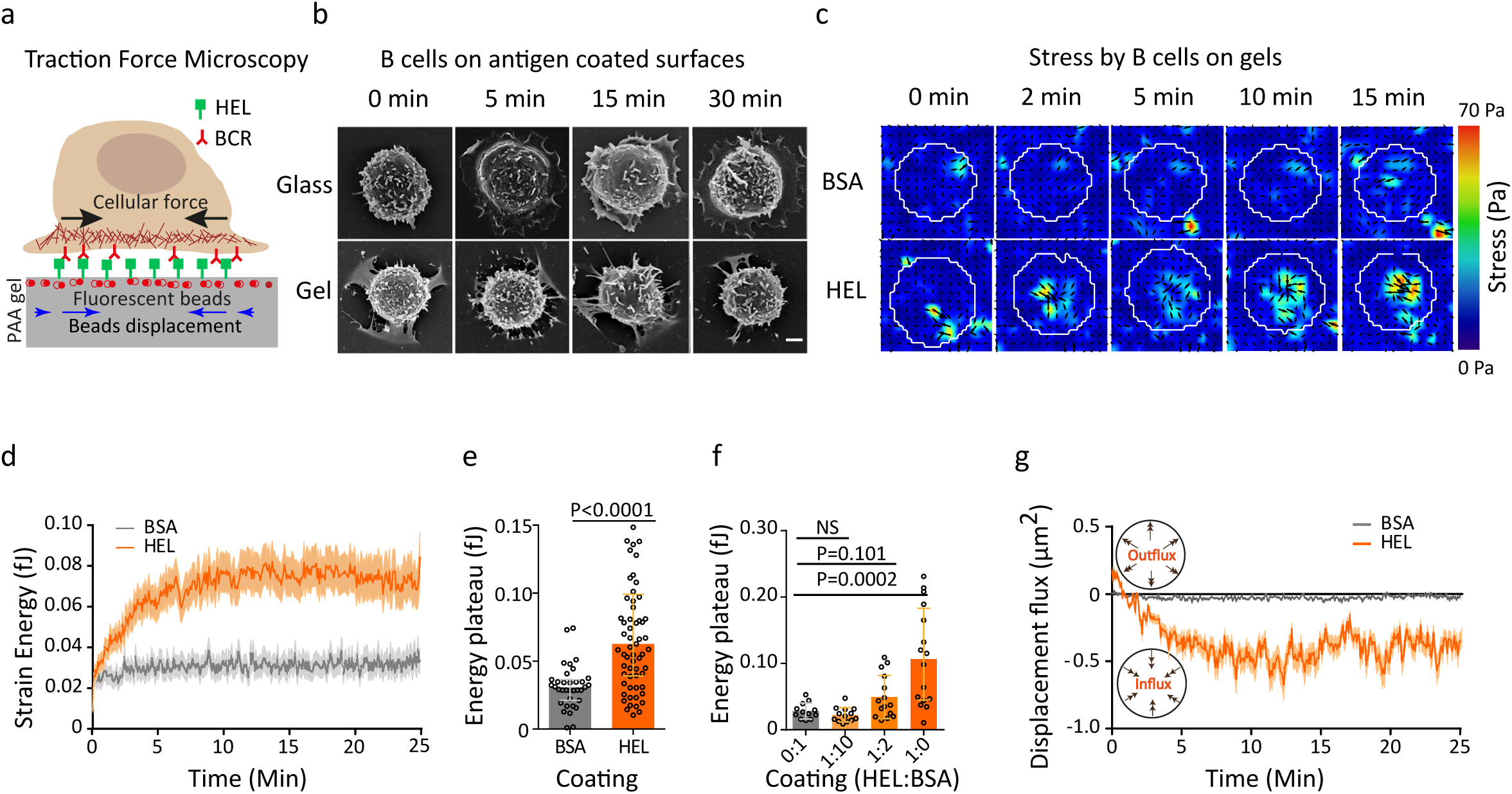
**B cells show antigen specific traction forces on PAA gels: (a)** Cartoon of traction force microscopy showing B cell plated on antigen coated polyacrylamide (PAA) gel containing fiducial markers. **(b)** Scanning electron microscopy of fixed B lymphocytes on HEL coated glass and PAA gels, scale bar is 2μm **(c)** Time-lapse color maps of stress for HEL and control BSA condition; shear stress can reach 70 Pa **(d)** Comparison of average strain energy profile for HEL and BSA conditions, error bars represent Mean±SEM (n=65 for HEL and n=35 for BSA, 5 independent experiments, 5 mice), acquisitions were started before the arrivals of the cells to capture the initial time of contact and all cells were aligned at time zero **(e)** Summary statistics of plateau of strain energy for HEL and BSA, error bar represents median±IQR (n=65 for HEL and n=35 for BSA, 5 independent experiments, 5 mice), Mann-Whitney test was performed for statistical analysis. **(f)** Concentration dependent increase in strain energy, error bars representing median±IQR (n = 10-15, 3 independent experiments, 3 mice), Mann Whitney test was performed for statistical analysis. **(g)** Displacement flux showing the direction of displacement over time in HEL and BSA condition (mean±SEM, n=65 for HEL and n=35 for BSA, 5 independent experiments, 5 mice).

Do these pulsatile contractile forces exerted on the substrate as a result of myosin II activity? To address this question, we generated conditional knockout mice in which *MYH9*, the gene coding for the main myosin II isoform expressed in lymphocytes (Immunological Genome Project, http://ww.immgen.org), was deleted in B cells **(Fig. 2a, Supp. Fig. 2a, Movie 3)** and expressed the HEL-specific BCR transgene. SEM analysis showed that B cells extracted from these animals did not show major morphological differences as compared to their wild-type counterpart **(Fig. 2b)**. In contrast, when measuring forces, we found that the absence of myosin II considerably decreased the contractile strain energy of the cells **(Fig. 2c,d,e)**. Still, measurable forces detected in both myosin II WT and KO cells were contractile, as shown by the negative (inward) displacement field fluxes **(Fig. 2f)**. Similar results were obtained when inhibiting myosin II with para-nitro-blebbistatin **(Supp. Fig. 2b)**. We conclude that the traction forces exerted at the B cell synapse depend on myosin II, as shown by both genetic deletion and pharmacological inhibition of the motor protein.

**Figure 2:**
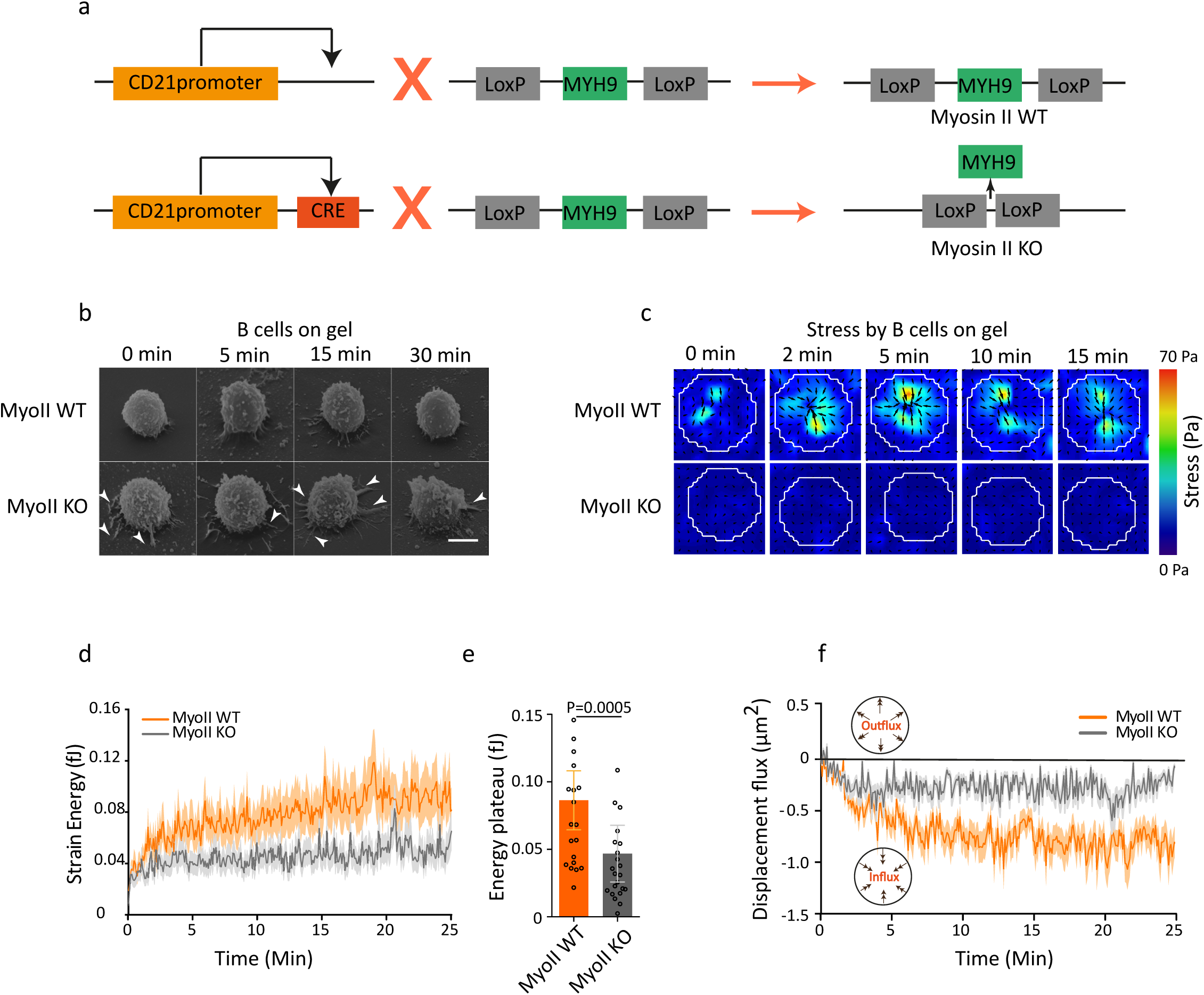
**Myosin II is essential for force generation by B cells: (a)** Genetic approach used to ablate Myosin IIA specifically in B cells: Myoll Flox mice are crossed with CRE+ mice under CD21 promoter. **(b)** Scanning electron microscopy of fixed B lymphocytes in Myosin IIA KO and WT condition, scale bar is 5μm **(c)** Timelapse images of stress colour maps for myosin II KO and WT condition, shear stress was of the order of 12 Pa **(d)** Average energy profile for myosin II KO and WT conditions, error bars represent Mean±SEM (n=23, 4 independent experiments, 4 mice). **(e)** Summary statistics of plateau of strain energy for myosin II KO and WT, error bars represent median±IQR (n=23, 4 independent experiments, 4 mice), Mann Whitney test was performed for statistical analysis. **(f)** Displacement flux showing the direction of displacement over time in myosin II KO and WT condition (n=23, 4 independent experiments, 4 mice).

An important hypothesis used to build the algorithm for force calculation in typical TFM experiments is that the displacement of cell-associated beads is accompanied by the displacement of its neighbours. However, we consistently observed that certain beads did not display movements parallel to the ones of their neighbours **(Fig. 3a, Movie 1)**. This apparently aberrant bead movement did not result from a modification of gel elasticity as the gel relaxed upon cell detachment (**Supp. Fig. 3a**). This suggests that these movements rather resulted from forces applied locally, possibly perpendicularly to the synaptic plan (which cannot be measured due to the limits of detection in z). We thus investigated the nature of these “point-like forces” by splitting the pool of beads into two groups based on *r*, the correlation between the directions of displacement vectors with its neighbours in a range of 1μm (**Fig. 3b**). For each frame, we classified the beads in two groups: *coordinated* (*r*>0.5) and *non-coordinated* (*r*<0.5). We found that all cells displayed both coordinated and non-coordinated movements in various proportions at each time point (**Supp. Fig. 3b**). Strikingly, these two types of movements were spatially segregated, as observed from average bead density maps and radial scans (**Fig. 3c**). The coordinated pool was located at the periphery of the synapse (~2-3μm from the centre), whereas the non-coordinated one was located at the centre of the synapse. Calculation of average coordinated displacements and relative stresses for each cell at each time point showed that shear stresses operated at the periphery and were organized centripetally. In contrast non-coordinated displacements were generally located at the centre of the cell-gel interface (**Fig. 3d**). Although the TFM algorithm underestimated point-like forces (**Supp. Mat.**), their distribution and relative value on average remained meaningful. Indeed, the number of beads moving in a coordinated manner increased with time, reaching a plateau about 2 to 3 min after initial contact (**Supp. Fig. 3c**). In contrast, the number of beads moving in a non-coordinated manner decreased in time. These results show that force patterning occurs in the first minutes after contact, during the rising phase of the strain energy. Dynamics analysis highlighted that coordinated forces organized in time as peaks that were correlated with isolated global contractions (**Fig. 3e**) and spectral analysis revealed a pseudo-period of 170±10s (median ± IQR) (**Fig. 3f and Supp. Fig. 3d**). We conclude that the forces at the B cell synapse present a specific spatio-temporal pattern with a peripheral, centripetal, pulsatile shear pool opposed to a central, point-like and disorganized one.

**Figure 3:**
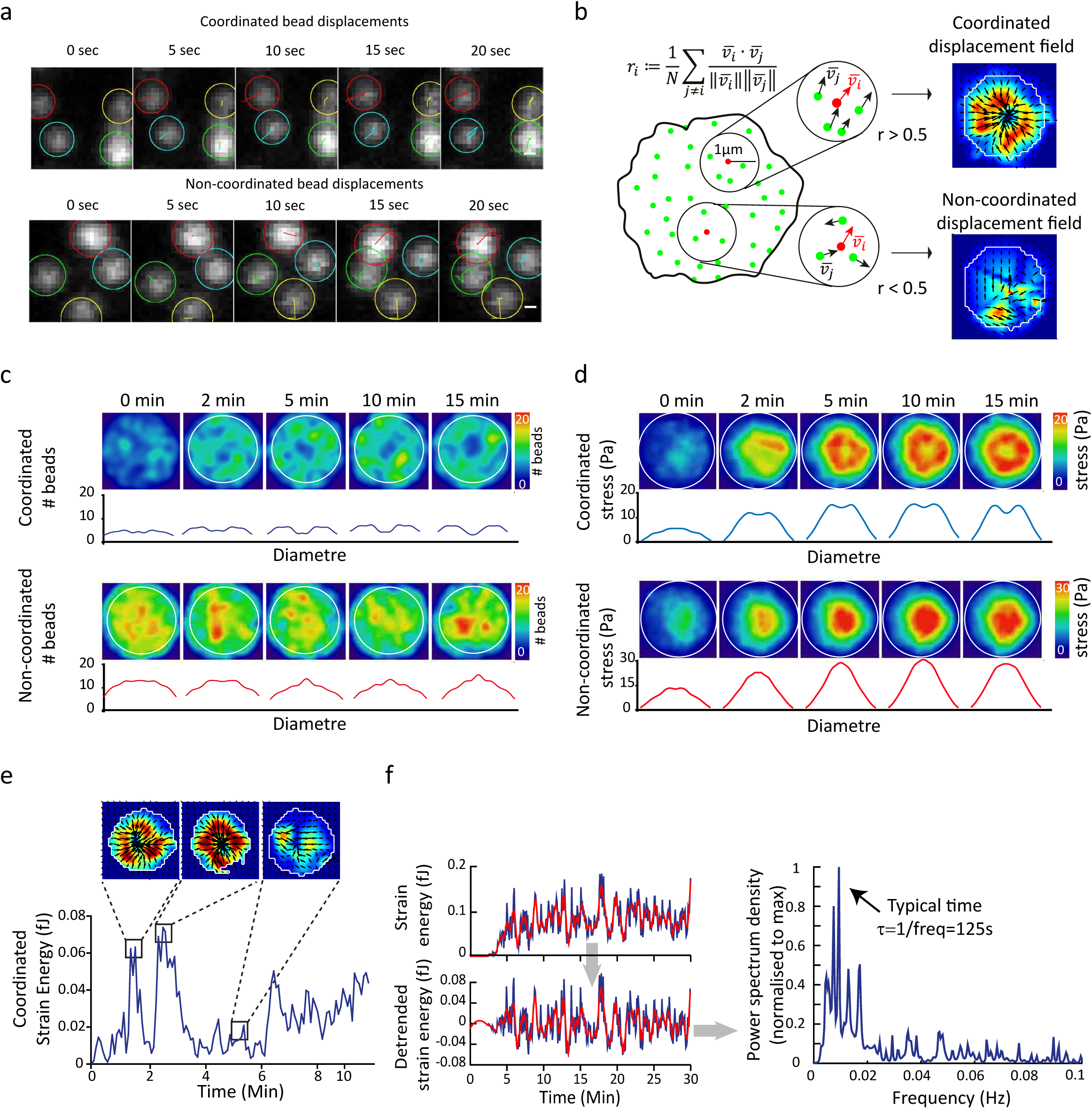
**Forces at the synapse exhibit two components: (a)** Time-lapse images of individual beads showing displacement tracks in coordinated and non-coordinated type of movements, scale bar is 2 μm. **(b)** Scheme showing the method used to separate the bead population in two pools, allowing extraction of coordinated and non-coordinated displacements. **(c)** Mean bead distribution in coordinated and non-coordinated pool of stress (n=100 cells). Below: radial profile of the density map obtained by average spatial distribution of the beads by resizing all cells and interpolating each bead with a Gaussian kernel. **(d)** Mean stress density map obtained by resizing and averaging at the indicated time point over all cells the individuals coordinated and non-coordinated stress maps. **(e)** Isolated peaks in coordinated strain energy correspond to peripheral ring of coordinated stress. **(f)** Extraction of the typical pulsation frequency from the coordinated contractile energy: from the time series of a coordinated energy signal (smoothed in red), we de-trended the signal and derived the power spectrum density that shows a maximum, hence a typical timescale for the pulsation.

How does myosin II contribute to each pool of forces? Monitoring the dynamics of myosin II-GFP at the cell-bead interface showed that the collective contractions described above corresponded to centripetal pulsations of myosin II, as shown by peaks in fluorescence intensity **(Fig. 4a, Movie 4)**. This observation became even more evident when the results were averaged over 40 different peaks **(Fig. 4b)**. Cross-correlation analysis showed that myosin II peaks preceded the shear strain energy ones by at least five seconds, suggesting that the motor is first recruited and then triggers global contractions. A drastically different result was obtained when analysing LifeAct-GFP dynamics at the cell-gel interface: the major actin pool observed corresponded to patches localized at the centre of the synapse (**Fig. 4c,d and Movie 5**). Noticeably, the regions where bead movements were non-coordinated were closely apposed to these actin patches (**Fig. 4e**), the two components being correlated over time, with no time lag (**Fig. 4f**). Are these actin patches responsible for the generation of the point-like forces observed at the centre of the synapse (**Fig. 4e**)? To test this hypothesis, we presented laterally pieces of antigen-coated gel to B cells. These experiments showed the presence of actin-rich protrusions that penetrated within the gel and were associated to bead movement **(Fig. 4g)**. The existence of these actin protrusions was further confirmed by cryo-electron microscopy **(Fig. 4h)**. Hence, the two pools of forces that are produced at the cell-gel interface are intrinsically different: the (shear) peripheral pool of coordinated forces shows pulsatile shear myosin II-associated centripetal contractions whereas the point-like central pool of non-coordinated forces is associated to dynamic actin-rich protrusions that form at the synapse centre. Interestingly, when measuring the coordinated and non-coordinated force components in the absence of myosin II, we found that they were both decreased, even though the effect was stronger on coordinated forces (**Fig. 4c**). Consistent with these findings, actin patches were lost in myosin II-inhibited cells (**Fig. 4c**), indicating that the activity of the motor protein is needed for the formation and/or maintenance of these actin protrusive structures. Altogether, these results suggest that force patterning at the B cell immune synapse results from contractility of the actomyosin cytoskeleton.

**Figure 4:**
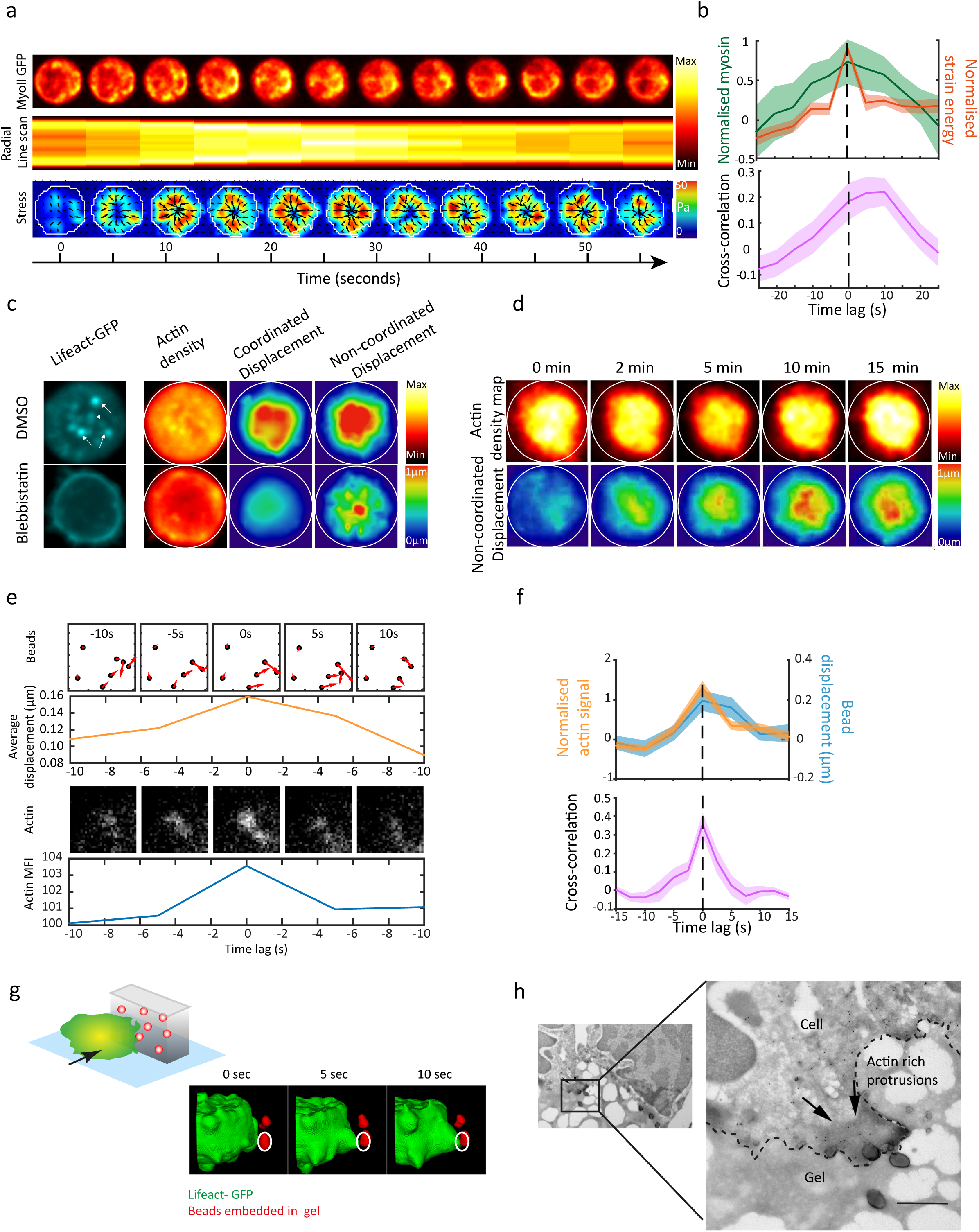
**Myosin II and actin dynamics: (a)** Time-lapse images showing dynamics of myosin II GFP, radial linescan (presented as a kymograph, time in x-axis) and corresponding stress map: from the linescan and the stress map a contraction peak is visible at times t=20-30s. **(b)** Average of different peaks and cross-correlation between strain energy and myosin II peak intensity: myosin peak precedes the energy peak of about 5-10 seconds (Mean±SEM, n=40 peaks, from 15 cells, 4 independent experiments). **(c)** Left, single cell showing actin patches in the centre of the cell which are lost in case of para-Nitroblebbistatin treated cells. Right, Mean actin distribution, mean coordinated and mean non-coordinated force density colour map in control and para-Nitroblebbistatin treated B cells (n=12). **(d)** Actin distribution over time correlates with non-coordinated force distribution (images are average projection of 12 cells over all time points). **(e)** Time-lapse images showing simultaneous appearance of an actin patch and non-coordinated bead displacements, arrowhead showing direction and magnitude of displacement; graphs below the images represent the respective signals integrated over a square of 3µm × 3µm. **(f)** Average of the signals quantified in (e) and average cross-correlation: both signals appear simultaneously and show a peak simultaneously with no lag (Mean±SEM, n=89, from 9 cells, 2 independent experiments) **(g)** Protrusions associated with bead movement (red) on antigen coated gels pieces presented laterally to B cell (green), 3D reconstruction. **(h)** Electron microscopy image showing actin protrusions through PAA gel (scale bar 0.2 µM).

These results prompted us investigating whether force patterning at the synapse impacts on the efficiency and localization of antigen extraction by B cells. For this, we recorded B cells plated on gels coated with fluorescently labelled antigen. Surprisingly, we observed that fluorescence was quenched when HEL was linked to the gel, being only detected upon HEL detachment (**Supp Fig. 4a**). This unexpected observation provided us with a robust system to measure HEL extraction at the B cell synapse and its dependency on force patterning. Time-lapse analysis showed a gradual detachment of antigen starting as soon as B cells contact the gel surface and slowing down ~3-5 min later **(Fig. 5a, Supp. Fig. 4b, Movie 6)**. This crossover time corresponded to the time at which the plateau was reached in the energy curve. Strikingly, the appearance of actin patches at the centre of the synapse coincided in space **(Supp. Fig. 4c and Movie 7)** and time with the appearance of HEL clusters (**Fig 5b**), indicating that actin protrusions are the sites where extraction of surface-tethered antigens occurs. Antigen detachment was impaired in the absence of myosin activity (**Fig. 5c**), in agreement with this motor protein being necessary for force production and formation of actin patches. Importantly, antigens were not only detached from the substrate, but also internalized within B cells, as shown by inside-out HEL staining **(Fig. 5d)** and cryoimmuno-electron microscopy **(Fig. 5e)**. These experiments further revealed the presence of myosin II on the cytosolic face of vesicles containing internalized antigens. Remarkably, when coating the gel with both fluorescent ovalbumin (OVA, not recognized by the BCR) and HEL, we found that only HEL was extracted **(Supp. Fig. 4d)**. This result shows that extraction exclusively concerns antigens that are bound to the BCR rather than entire pieces of the gel, indicating that antigen extraction at the B cell synapse (at last in this case) is distinct from the trogocytic process previously reported at T and B synapse ([20], [21]). We conclude that B cells use myosin II to form actin patches where surface-tethered antigens are internalized. However, myosin II is not recruited to these patches.

**Figure 5:**
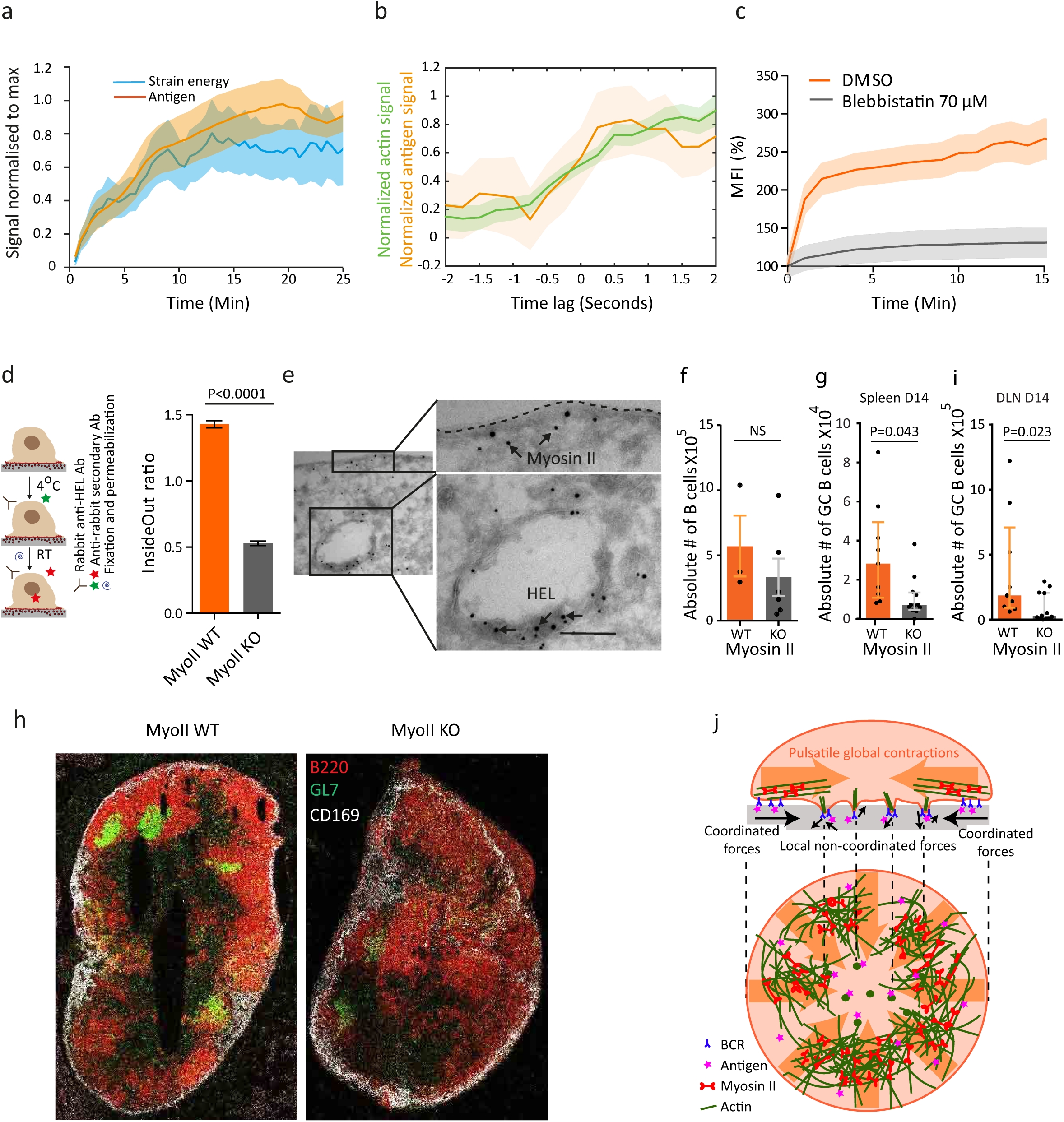
**Antigen Internalization**: **(a)** Strain energy and antigen gathering (Mean±SEM, n=15, 5 independent experiments); signals are normalized to the maximum to highlight similarity in the kinetics. **(b)** Signals of the fluorescent HEL and Lifeact GFP patches integrated over a square of 3µm × 3µm: both signals appear simultaneously (Mean±SEM, n=21, from 6 cells). **(c)** Antigen gathering over time in control and para-Nitroblebbistatin treated cells (error bars show Mean±SEM, n=15 DMSO, n=9 Blebbistatin). **(d)** Scheme of the inside-out experiment and relative quantification: ratio of internalized (inside) and not internalized (outside) antigen in myosin II KO and WT B cells, error bars showing Mean±SEM (n=29 Myosin II WT, n=21, Myosin II KO, 3 independent experiments, T-test was performed for statistical analysis). **(e)** Scanning electron microscopy images showing internalized antigens and its proximity to myosin II (scale bar 0.2µM). **In vivo observations: (f)** Absolute number of CD19 positive B cells in myosin II WT and KO mice (each dot represents one mice, 2 independent experiments, error bars represents mean±SEM, Mann Whitney test was performed for statistical analysis). **(g)** Absolute number of germinal centre B cells in spleen and **(i)** draining lymph node in myosin II WT and KO beads immunized mice (each dot represents one mouse, 2 independent experiments, error bars represent median±IQR, Mann Whitney test was performed for statistical analysis). **(h)** Histology image of draining lymph node from immunized mice showing B cells (B220), germinal centres (GL7) and sub-capsular sinus macrophages (CD169); images highlight scattered germinal centre B cells in myosin II KO mice. **(j) Model:** myosin II driven global peripheral contractions (shear coordinated forces) allow the endocytic machinery to build up in the centre where antigen is extracted through actin protrusions associated to the generation of localised forces.

We finally evaluated the relevance of these findings *in vivo* by comparing B cell responses in WT mice and mice whose CD21^+^ B cells were KO for the myosin IIA gene. We found that the number of B cells in lymph nodes did not differ significantly between WT and myosin II KO mice **(Fig. 5f)**. However, upon immunization with surface-tethered antigens, the formation of germinal centres was impaired in myosin II KO mice **(Fig. 5g,h and Supp. Fig. 4e).** Histological analyses showed that germinal centres were not only decreased in number but also disorganized in the absence of the motor protein **(Fig. 5i)**. We conclude that myosin II is required for B lymphocytes to mount efficient immune response responses *in vivo*.

In summary, we here found that stress at the synapse is patterned into two components: a shear peripheral component, mainly associated to pulsatile acto-myosin contractions, and a protrusive normal component, concentrated at the centre of the synapse and associated to transient actin protrusions where antigens are extracted. Peripheral pulsatile dynamics of actomyosin contractions might be needed for antigen internalization to occur at the centre of the synapse, most likely by the formation of actin patches. Indeed, little HEL detachment was observed at the periphery of the cell, suggesting that shear coordinated forces generated by myosin II pulsatile contractions do not directly contribute to antigen extraction. Interestingly, studies on synapse organization on lipid bilayers show that actin organizes in concentric structures crossed by radial fibres [22] through continuous centripetal flow [12], [14]. In B lymphocytes interacting with antigen-coated gels, we did not observe such actin flows but collective pulsatile actomyosin contractions, similar in features and timescales to the ones described in several other systems [23]-[26]. Are these pulsatile contractions *sufficient* to form actin patches and thus localize antigen extraction at the centre of the synapse? The model presented in (**Fig. 5j**) suggests that they are. This model considers molecules attached to the membrane as diffusive objects advected by an intermittent flow: their diffusion time (i.e. the one necessary to destroy patterns) is relatively long, larger than the typical period of the pulsations observed (see **Supp. Mat.**). We show that this condition is sufficient to generate a radial gradient. These patterns might explain the formation of particular structures such as actin protrusions at the centre of the synapse, as it has been shown that an active gel coupled to the cell membrane forms aster like patches [27], [28].

Myosin II-dependent pulling was proposed to protect B cells from internalizing low-affinity antigens by mechanically disrupting the lose bonds they form with the BCR. However, mechanical affinity discrimination is the result of tiny differences in energetic barrier (see **Supp. Mat.**) and is likely to be disturbed by the external noise (lymphatic flows, muscle contractions, cell motility, down to molecular thermal noise). In this context, the peripheral stress may help sealing the synapse and mechanically isolate it from external noise. Similar scenario might apply to the T cell synapse and would deserve further investigations as myosin II also plays an important role in the forces exerted by these cells [29]-[33].

Spatio-temporal force patterning was first highlighted in the context of tissues [34], cell adhesion to substrate [35]-[37] and cell motility [25]. Our study shows that it might be a more general and basic feature of cell-cell interfaces where the engagement of surface receptors leads to both juxtacrine signalling and ligand endocytosis. We found that myosin II intervenes in this process as a global and local master organizer of forces and actin distribution, and thus as a key actor of endocytosis which is a key process of adaptive immunity. We anticipate that this study should set the ground for future work aimed at exploring force patterning in additional cell-to-cell communication models.

## Acknowledgements

We acknowledge Y. Bellaiche, G. Cappello, M. Cosentino-Lagomarsino, C. Hivroz, J. Husson, I. Lavi, M. Leoni, O. Malbec, H. Moreau, M. Piel, J. Pineau and P.H. Puech for discussions, advices about experiments and critical reading of the paper. We acknowledge the Nikon Imaging Center@CNRS-Institut Curie and PICT-IBiSA, Institut Curie, Paris, member of the France-BioImaging national research infrastructure, for support in image acquisition and the Curie Mouse Facility. AK was supported by Paris Descartes PhD fellowship. This project was funded by grants to PP(ANR-10-JCJC-1504-Immuphy), AMLD (ANR-PoLyBex-12-BSV3-0014-001, ERC-Strapacemi-GA 243103), MCIK (Swedish Research Council).

## Author contributions

A.K. performed and analysed most experiments, prepared figures and participate in writing the first draft. P.J.S. helped with the experimental setup and cell manipulation. M.M. performed image analysis. D.L. performed immuno-staining and SEM experiments, M.S.-R. performed TEM experiments. R.V. build the analytical model and contributed to analytical arguments. K.H. performed preliminary TFM analysis. V.F.B. performed and analysed *in vivo* experiments. M.C.I.K. supervised the *in vivo* experiments. M.B. provided TFM expertise and analytical tools. A-M.L.D. designed experiments, wrote the manuscript and supervised the whole research. P.P. designed experiments, performed formal and data analysis, prepared figures, wrote the manuscript and supervised the whole research.

---

**supp 1:**
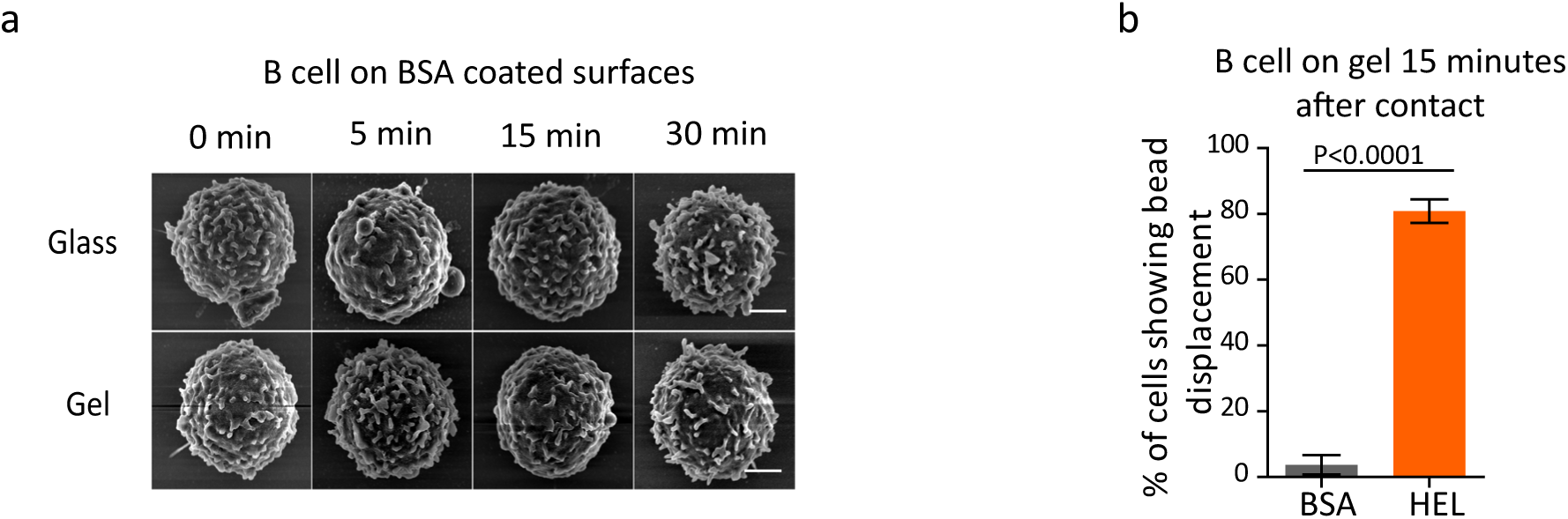
**(a)** Scanning electron microscopy of fixed B cells on BSA coated glass and polyacrylamide gels, scale bar is 2μm. **(b)** Percentage of B cells showing bead displacement in HEL and control BSA condition (n=10 mice), error bars represents mean±SEM, Mann Whitney test was performed for statistical analysis.

**supp 2:**
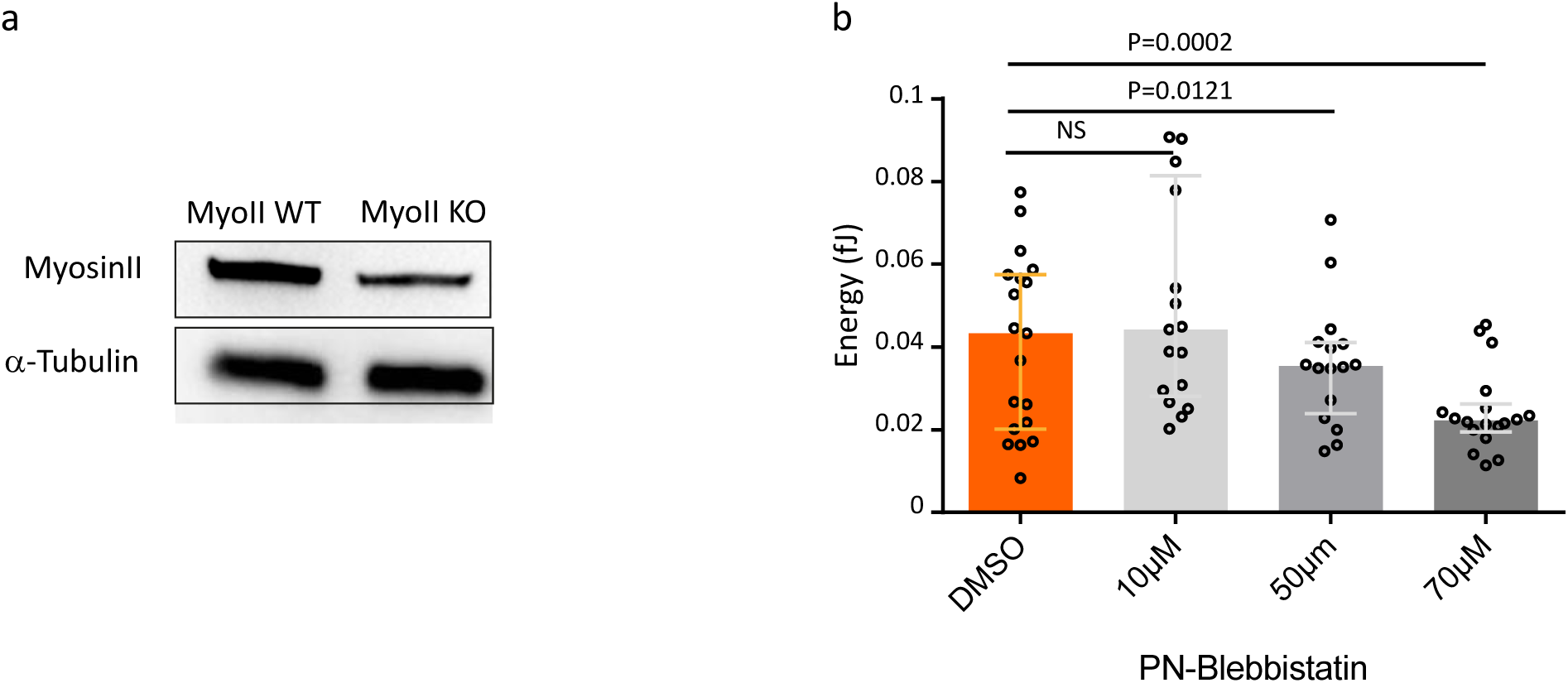
**(a)** Western blot analysis of the efficiency of Myosin II knock out, α-tubulin was used as loading control (example selected out of n=4 experiments, all showing the same tendency). **(b)** Concentration dependent decrease in strain energy of para-Nitroblebbistatin treated B cells (error bars representing median±IQR, n=10-15, 3 independent experiments, 3 mice, Mann Whitney test was performed for statistical analysis).

**supp 3:**
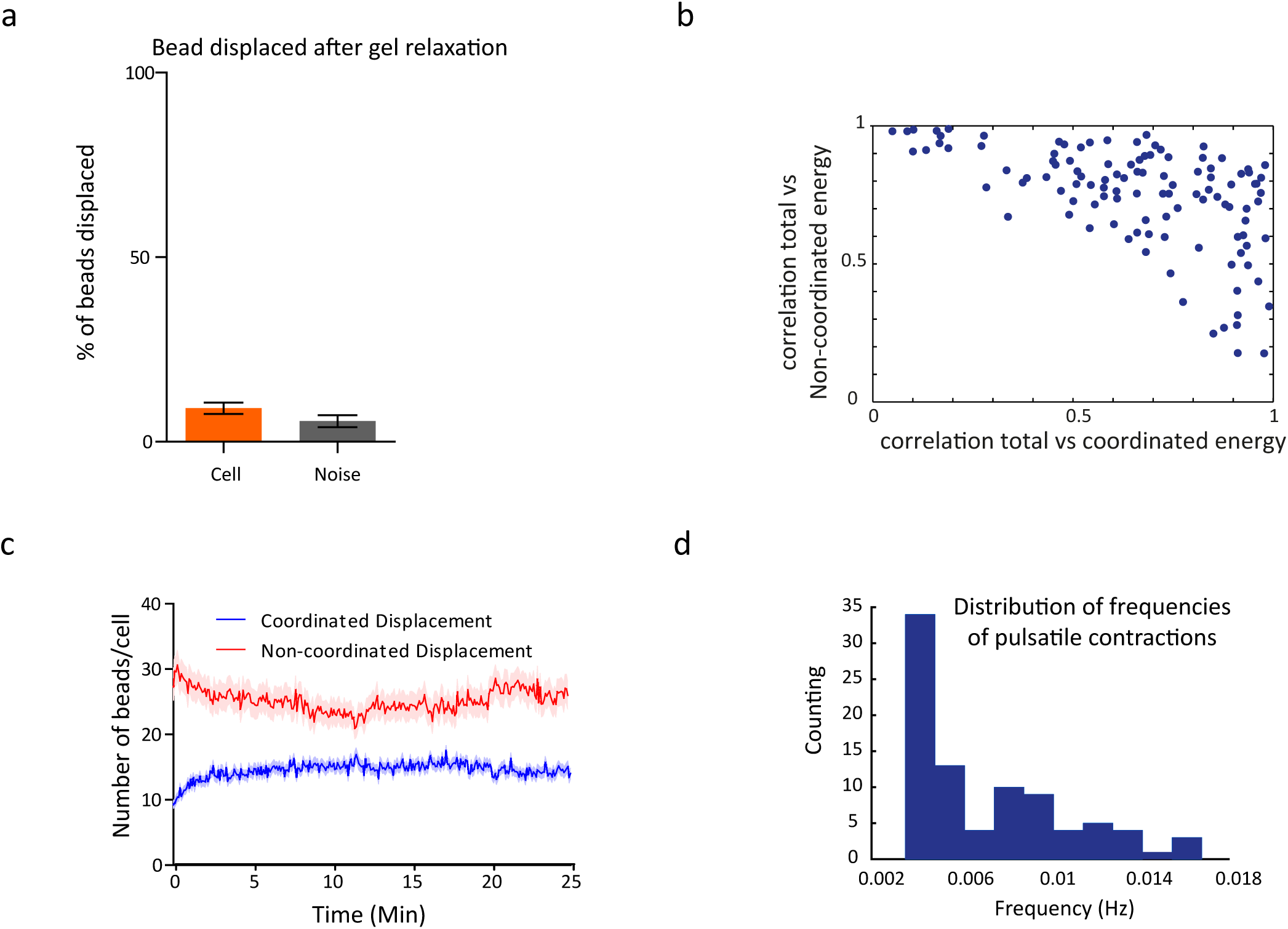
**(a)** Non-coordinated displacements are not resulting from gel deterioration: percentage of bead displaced on removing the cells after the experiments, Mean±SEM (n=15), Mann Whitney test was performed for statistical analysis. **(b)** Distribution of each cell in the profile shows correlation of total forces to coordinated force and non-coordinated forces respectively. **(c)** Number of beads per cell over time in coordinated and non-coordinated type of forces. Error bars represent Mean±SEM. **(d)** Histogram of the spectra maxima in frequency returning a median typical time of 170s for the period of myosin contractions (n=45).

**supp 4:**
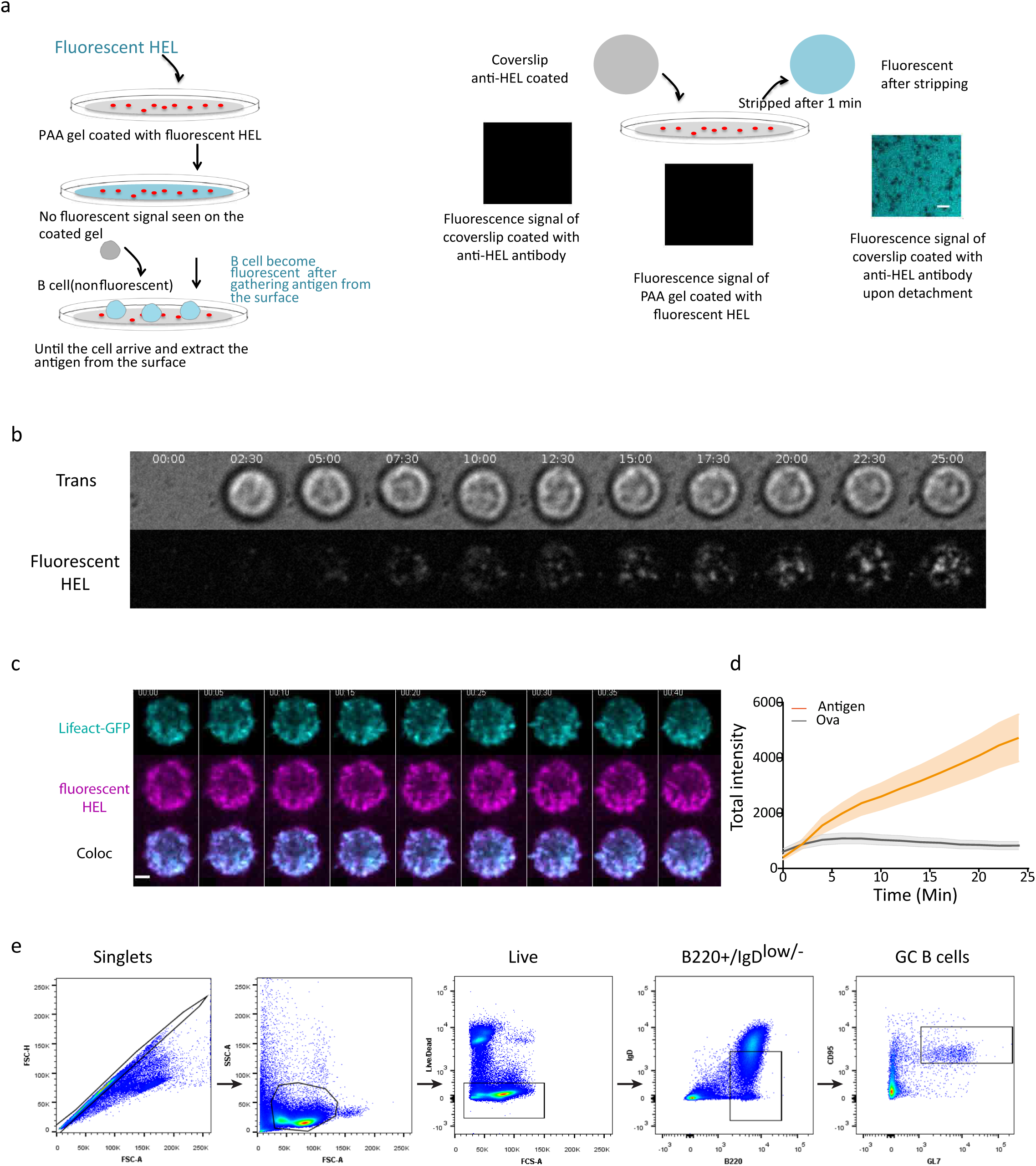
**(a)** Scheme showing stripping experiments: anti-HEL coated coverslip (non fluorescent) was put 1 minute in contact with fluorescent HEL coated gel (non fluorescent) and successively stripped; the coverslip becomes fluorescent (scale bar is 2 μm). **(b)** Antigen extraction begins at the centre of the synapse as soon as the cell touches the surface. **(c)** Time lapse image of fluorescent HEL and Lifeact-GFP showing colocalization of HEL and actin puncta. **(d)** Antigen extraction is specific: fluorescent HEL intensity upon B cell contact compared to fluorescent Ova, error bars represent Mean±SEM (n=67 Antigen, n=67 Ova, 2 independent experiments). **(e)** Gating strategy for selection of the germinal centre B cell: these cells are selected as B220+/IgD low/CD95+/GL7+.

## Movies

Movie 1: beads displacements in antigen coated gel showing global pulsatile contractions. One image every 5s (0.2fps), scale bar 2μm.

Movie 2: Stress maps (and quiver plot) generated by B cells on HEL and BSA coated substrates. Movies were generated after analysis of the beads displacement using TFM inversion algorithm. One image every 5s (0.2 fps).

Movie 3: Stress maps (and quiver plot) generated by myosin II WT and KO B cells on HEL coated substrates. Movies were generated after analysis of the beads displacement using TFM inversion algorithm. One image every 5s (0.2fps)

Movie 4: myosin II-GFP contractions. From left to right: transmission, stress map (and quiver) and Myosin II GFP signal. Two global contractions are highlighted. One image every 5s (0.2fps).

Movie 5: lifeact-GFP patches. From left to right: transmission, fluorescent beads and Lifeact-GFP. One image every 5s (0.2fps), scale bar 2μm.

Movie 6: fluorescent antigen extracted form the gel. Left transmission, right: fluorescent HEL (appearing of fluorecncens as long as it is extracted. One image every 5s (0.2fps), scale bar 2μm.

Movie 7: lifeact-GFP and fluorescent HEL (AF-532). From left to right: Actin (cyan), HEL (magenta), colocalization (white). One image every 0.5s (2fps), scale bar 2μm.

## Materials and methods

**Mice and cells**: Mice with a conditional deletion of myosin II in B cells were generated by backcrossing mice carrying a floxed myosin II allele (*MyosinIIflox/flox*) [1] with mice expressing the Cre recombinase under the control of the CD21 promoter (*CD21 cre*^+/−^). Mice expressing the hen egg lysozyme (HEL)-specific MD4 receptor were also crossed with mice carrying a floxed myosin II allele. Mice were crossed at an age of eight to 10 weeks, and Cre^−^ littermates were used as WT controls. The transgenic MD4, Lifeact GFP and myosin II-GFP mouse lines have been described elsewhere [2], [3]. This resulted in all the desired genetic combinations being obtained in the C57BL/B6 background, and the corresponding breeding controls were systematically used. The experiments were performed on 8 to 10 week-old male or female mice. Animal care conformed strictly to European and French national regulations for the protection of vertebrate animals used for experimental and other scientific purposes (Directive 2010/63; French Decree 2013-118). Immunization experiments carried out at the Karolinska Institute were performed according to local ethical committee guidelines (N11/13). Mature spleen B cells were purified with the MACS kit (130-090-862). B cells were cultured ([4]) in CLICK medium (RPMI 1640—GlutaMax-I supplemented with 10% fetal calf serum, 1% penicillin-streptomycin, 0.1% β-mercaptoethanol and 2% sodium pyruvate.

**Antibodies and reagents:** The following reagents were used: 100 μg/ml HEL (Sigma), 100 μg/ml BSA (Euromedex), 40% polyacrylamide (Biorad), 2% bis-poylacrylamide (Biorad), 3-aminopropyltrimethoxysilane (Sigma), 0.2 μm Alexa647 Fluospheres (Thermo Fisher, F8807), Sigmacote (Sigmacote), ammonium persulfate (Sigma), TEMED (ICN Biomedicals), Sulfo-SANPAH (Thermo Fisher), and the Alexa555 protein labeling kit (A30007, Molecular Probes). For drug treatments, we used para-nitro blebbistatin (Optopharma). The following antibodies were used: rabbit anti-HEL (Abcam, 1/100), human anti-GFP (Institut Curie, 1/200), Alexa Fluor 647-conjugated anti-phalloidin (Thermo Fisher, 1/200), anti-myosin IIA heavy chain (Covance, 1/500), Alexa Fluor 488-conjugated goat anti rabbit IgG(Life Technologies, 1/200), Alexa Fluor 488-conjugated goat anti-human IgG (Life Technologies, 1/200).

**Inhibitors:** Cells were incubated with 70 μM para-nitro blebbistatin (Optopharma) for 40 minutes at 37°C in RPMI media before the experiments, unless otherwise stated.

**Live cell microscopy:** Fluorodishes containing gels were placed under a microscope focused on the plane of beads. In total, 1×10^5^ B cells were added to the plate (time 0), and images were acquired over time. Images were acquired at 37°C, under an atmosphere containing 5% CO_2_, with an inverted spinning disk confocal microscope (Roper/Nikon) equipped with a 60X (1.4 numerical aperture (NA)) oil immersion objective and a CoolSNAP HQ2 camera. Images were acquired in the synaptic plane, at five-second intervals.

**Preparation of polyacrylamide gel substrates:** PAA were produced in 35-mm FD35 fluorodishes (World Precision Instruments, Inc). These dishes were first treated by UV irradiation for 2 minutes, and then with 3-aminopropyltrimethoxysilane (APTMS) for 5 minutes. The dishes were washed thoroughly in distilled water before preparation of the polyacrylamide gels. Hydrophobic coverslips were prepared by incubation in Sigmacote for 3 minutes, followed by thorough washing and drying. A 500 Pa gel was prepared by diluting 40% polyacrylamide and 2% bis-acrylamide solutions to obtain stock solutions of 12% acrylamide/0.1% bis-acrylamide. We sonicated 167 μl of this solution with 10% of 0.2 μm fluorescent beads, and then added 0.2 μL of TEMED and 10% ammonium persulfate and mixed thoroughly, to initiate polymerization. A volume of 9 μL of the polyacrylamide mixture was immediately pipetted onto the surface of the Fluorodish and a Sigmacote activated coverslip was carefully placed on top. Fluorodishes were immediately inverted, to bring the beads to the surface of the gel. Polymerization was completed in 45 min and the top coverslip was then slowly peeled off and immediately immersed in PBS. Sulfo-SANPAH was used to crosslink antigen to the surface of the gel. It is a surface functionalizing reagent with an amine-binding group that binds to the antigen (HEL, IgG, and IgM) and a photoactivable azide group that binds to the substrate upon activation under UV giving homogeneous distribution of antigen on the surface of PAA gels. We pipetted 150 μl of 0.5 mg/ml sulfo-SANPAH onto the surface and placed the gel under UV light for 2 minutes. The gels were then washed with PBS and the process repeated. The gel was washed thoroughly with PBS and coated with 100 μg/ml antigen, by overnight incubation at 4°C.

**Characterization of the polyacrylamide gels:** The Young’s modulus of PAA gel was measured by bead indentation and calculated using a Hertz model for an elastic substrate with finite thickness [5]. Glass beads of 0.25 mm radius were deposed on the gel and their indentation was measured using confocal stacks. Gel height was determined by focussing on the bottom and top of the gel. The force inserted in the Hertz formula was computed theoretically as the weight of the glass bead (density=2.2kg/m^3^ and radius 0.25mm) minus the buoyancy in water. PAA gels in our system remains in the range of 400-650 Pa.

**Quantification of amount of antigen on PAA gel and glass:** To be sure that the difference of spreading on gel and on glass was not due to the amount of antigen coated on different substrates (it is harder to coat gels with protein due to the inherent hydrophobicity of the PAA). We inferred the amount of antigen required for coating the glass with an equivalent concentration of antigen on gel (100μg/ml) by taking images at different concentration on glass and comparing with the fluorescent intensity obtained on the gel of 100μg/ml. Respective glasses and gels were coated with HEL overnight at 4°C and later stained by using rabbit anti-HEL primary antibody at 37°C, eventually staining with anti-rabbit alexa-488 secondary antibody. Images were acquired using laser scanning microscope (Leica) with a 40X 1.4 NA oil immersion objective with 5% 488 laser. Mean fluorescence intensity at different point follows a logarithmic curve that suggests the equivalent concentration on glass is 0.14 μg/ml.

**Immunofluorescence:** B cells plated on polyacrylamide gels were fixed by incubation in 4% PFA for 10 min at room temperature, and PFA was quenched by incubation in PBS plus 1 mM glycine for 10 min. Fixed cells were incubated with primary antibodies in PBS plus 0.2% BSA and 0.05% saponin, overnight at 4°C. The cells were washed and incubated with secondary antibodies for 1 hour at room temperature. Immunofluorescence images were acquired on a laser scanning microscope (Leica SP8, software LAS X) with a 40× 1.4 NA oil immersion objective.

**Inside-out immunofluorescence:** B cells were plated on polyacrylamide gels and incubated for 15 min at 37°C. The cells were then transferred to 4°C and Fc receptors were blocked for 10 min using Fc blocker (BD, 1/200). The cells were washed with PBS and incubated with rabbit anti-HEL antibody at 4°C for one hour and then with Alexa Fluor 488-conjugated anti-rabbit IgG secondary antibody for one hour. The cells were moved to room temperature, fixed by incubation with 4% PFA for 10 minutes and permeablized by incubation with PBS plus 0.2% BSA and 0.05% saponin. The cells were then incubated with rabbit anti-HEL antibodies for one hour, washed with PBS-BSA-SAPONIN, and incubated with the Alexa 546-conjugated anti-rabbit IgG secondary antibody for one hour at room temperature. The cells were washed several times in PBS and then used for imaging. Images were acquired using laser scanning microscope (Leica) with a 40X 1.4 NA oil immersion objective.

**SEM Imaging:** Cells were fixed overnight at 4°C in 2.5% glutaraldehyde in phosphate buffer on 0.2μg/ml HEL coated on glass and 100μg/ml HEL coated on PAA gel. They were dehydrated in a graded series of ethanol solutions, then dried by the CO_2_ critical-point method, with an EM CPD300 (Leica microsystems). Samples were mounted on an aluminum stub with silver lacquer and sputter-coated with a 5 nm layer of platinum, with an EM ACE600 (Leica Microsystems). Images were acquired with a GeminiSEM 500 (Zeiss).

**Electron microcopy:** Immunoelectron microscopy was performed by the Tokuyasu method (Slot & Geuze, 2007). Double-immunogold labeling was performed on ultrathin cryosections with protein A-gold conjugates (PAG) (Utrecht University, The Netherlands). Electron micrographs were acquired on a Tecnai Spirit electron microscope (FEI, Eindhoven, The Netherlands) equipped with a Quemesa (SIS) 4k CCD camera (EMSIS GmbH, Münster, Germany).

**Western blotting:** B cells were lysed at 4⍛C in lysis buffer (10 mM Tris HCL pH 7.4, 150 mM NaCl, 0.5% NP40). Cell lysates were loaded onto mini-PROTEAN TGX SDS-PAGE gels, which were run at 200 volts and 65 mA. The bands on the gel were transferred onto Polyvinylidene fluoride (PVDF) membranes (Trans-Blot Turbo Transfer). Membranes were blocked with 5% BSA in 1xTBS (Tris buffered saline)-0.05% Tween-20 and incubated overnight at 4°C with primary antibodies and then for 60 min with secondary antibodies. Western blots were developed with Clarity Western ECL substrate, and chemiluminescence was detected with a ChemiDoc imager (all from BioRad).

**Density map analysis:** On movie reconstruction, individual cells were cropped with ImageJ software. For signal mapping, the images obtained for each individual cell were aligned in a single column. Cell size normalization was applied to each time point, based on mean cell size, with background subtraction. We obtained a mean behaviour for each cell, by projecting every time point onto the average. The mean behaviour of the population was then determined, by projecting the mean signal of every individual cell at a given time point. This procedure was performed with a custom-designed ImageJ-compatible macro. A similar procedure was used to map stresses, except that the real stress value was used, without normalization. For bead density analysis, we smoothed positions with a two-dimensional Gaussian kernel of radius 3 pixels to obtain a density map, as described by [6]. These last two analyses were performed in Matlab.

**Myosin and energy peak analysis:** maxima of the coordinated energy were isolated manually and a sequence of 11 frames around each maximum isolated and aligned to the maximum. The average Myosin II-GFP fluorescence was integrated in the area of the cell and aligned blindly following the energy sequence alignment. The pieces of signals were offset to zero and normalised to the maximum, averaged and plotted. For the correlation analysis the signals were cross-correlated and the average cross-correlation plotted.

**Actin patch and displacement analysis:** actin patches were isolated manually and the signal integrated in a square of 2×2 μm. In time sequence of 11 frames were considered separately. The signal of the displacement was computed as average absolute length of the displacement vector of the non-coordinated beads population in the same square used for the actin signal. Each sequence of actin signal was offset to zeros and aligned according to the maximum in fluorescence. The displacement signal was aligned blindly following actin ones. The pieces of signals were averaged and plotted. For the correlation analysis the signals were cross-correlated and the average cross-correlation plotted.

**Actin and HEL patches analysis:** similar analysis of the antigen patches but at hight acquisition rate, in a smaller sequence and in a smaller window were performed to show the simulatenous apparence of HEL patches. The only difference is that in this case the HEL signal was aligned first to the point of appearance and the actin signal was blindly translated.

**Antigen-stripping experiments:** Round glass coverslips were coated with anti-HEL antibody by overnight incubation. The coverslips were washed with PBS and imaged to obtain the control image. They were then placed on the antigen-coated polyacrylamide gel for 30 seconds to 1 minute. They were stripped off the surface of the gel and imaged as soon as possible using laser scanning microscope (Leica) with a 40× 1.4 NA oil immersion objective. Fluorescence on these images indicated the presence of detached antigen. Absence of fluorescence in stripped gel suggests that quenching is not due to the presence of many layer of antigen.

**Traction force Microscopy: energy and flux:** The traction force algorithm was based on that used by [7] and modified by [8]. Force reconstruction was conducted with the assumption that the substrate is a linear elastic half space, using Fourier Transform Traction Cytometry with Tikhonov regularization (regularization parameter was set to 5×10^−19^). The position of the beads in reference image and deformed one was measured using MTT algorithm [9]. The problem of calculating the stress field from the displacement is solved in Fourier space then inverted back to real space. The final stress field is obtained on a grid with 0.432 μm spacing (4 pixels). All calculations and image processing were performed in Matlab.

Given the size of B cells, the density of beads, the magnitude of displacement, some parameters needed optimization for the analysis. In particular for the detection algorithm (MTT): search window size (5 pixels), particle radius (2.5 pixels) and maximum distance for nearest neighbour (4 pixels). Pixel size of spinning disk confocal microscope is 108 nm (we occasionally used another setup with pixel size 160 nm, but the parameters did not need adjustment). Same parameters were applied for noise detection by measuring force in a non-stressed area not too far from the cell. A quality check of the TFM algorithm is given by the "non equilibrated forces", i.e. by the ratio of the sum of forces vectors (which should be zero) to the sum of magnitude of the forces. Lower ratio signifies higher quality of the analysis. We checked that all analysed data this ratio was below 0.15. Further calculations based on the output of the algorithm were performed to extract the total strain energy (defined as the sum over the entire cell area of the scalar product force by displacement). Fluxes were calculated by standard vector analysis (Green's theorem): the flux is the integral over the cell area of the divergence of the 2D field (displacement or stresses). An outward flux represents pushing forces and inward flux represents pulling forces.

Note that even if in theory the forces are supposed to be zero outside the cell, we decided not to introduce this constrain to avoid border effects. However when we compute energy and fluxes, we use the mask of the cell extracted by using an ImageJ custom made macro. The mask was increased by 10% (dilation of the binary image using Matlab morphological tools) to avoid excessive cropping of the force/displacement field.

**TFM algorithm to analyze coordinated and non-coordinated forces:** we determined whether a bead belongs to the coordinated or non-coordinated group, by calculating the mean correlation between the displacement vector associated with the bead and its nearest neighbours (within 1 μm range). Beads with a correlation coefficient below 0.5 were considered to belong to the non-coordinated pool. Note that we define a correlation that does not depend on the magnitude of the displacement vectors but only of their relative orientation. This implies that beads moving a little or not at all have low correlations coefficient and build up the non-coordinated pool. The scatter plot **(Supp. Fig. 3b)** based on the correlation was generated based on the correlation coefficient between the entire time series for total energy and the entire time series for energy associated with coordinated/non-coordinated beads, respectively, for the *x* and *y* axis.

**Spectral analysis:** To extract a typical time-scale of the collective pulsatile dynamics (Figure 3f) the coordinated energy was first de-trended subtracting the background obtained smoothing the original signal with Savitsky-Golay filter (with a window of 500s). The power spectrum was then computed on the detrended signal using maximum entropy algorithm (Matlab). The maximum was selected above frequencies of 1/500 Hz (to avoid including effect of the smoothing).

**FACS antibodies:** Cells were blocked with rat anti-mouse CD16/CD32 (BioLegend) and stained with: LIVE/DEAD™ Fixable Aqua Dead Cell (ThermoFischer), PerCpCy5.5 Rat anti-mouse IgD (BioLegend), Pacific Blue Rat anti-mouse B220 (BioLegend), PE Cy7 Hamster anti-mouse CD95 (BD Biosciences), PE Rat anti-mouse T and B cell activation antigen (BD Biosciences). Samples were attained on BD LSRFortessa X20 and analyzed using FlowJo software.

**Mice immunization:** HEL (Sigma)-OVA (Sigma) coated beads used in immunization experiments were prepared as follows: 7.5 μg of biotinylated HEL + 7.5 μg of biotinylated OVA 647 were incubated overnight at 4°C with 10^7^ streptavidin coated 200nm beads (Sigma) in 500μl, washed 4 times and re-suspended in PBS-BSA 1% at a concentration of 80×10^6^ beads/μl. Mice were injected subcutaneously in the left flank with 50μl beads in Alum (ThermoScientific) in a ratio 1:1 (mice received 4×10^9^ beads or 3μg HEL + 3μg OVA). Draining (inguinal) lymph nodes were collected on day 14.

**Lymph node immunofluorescence:** 8 μm thick lymphnode sections were blocked for 30 minutes with 5% goat serum (DakoCytomation) in PBS. GL7 antibody was incubated overnight at 4°C, followed by washing with PBS and staining of the remaining directly conjugated antibodies for 1h at room temperature. The following antibodies were used: Alexa Fluor 488-conjugated rat-anti-mouse T and B cell activation antigen (BioLegend) and PE-conjugated rat-anti-mouse B220 (BioLegend), Alexa Fluor 647-conjugated anti-mouse CD169 (BioLegend). Afterwards, the tissue sections were washed with PBS and mounted with Prolong Diamond mounting medium (Invitrogen). Images were collected using a confocal microscope (Zeiss LSM880) and analyzed using ImageJ software.

**Statistics:** All graphs and statistical analyses were performed with GraphPad Prism 5 (GraphPad Software) and MATLAB. In most cases non-paramteric test (Mann-Whitney) test was used to determine statistical significance unless otherwise stated. Bar graphs show the median ± interquartile range (IQR) or mean± standard error mean (SEM). Graphs representing strain energy and displacement flux were aligned to start at time zero, dot plots of strain energy show the average of each cell at the plateau.

**Code availability:** TFM analysis codes (Matlab) and image quantification tools (ImageJ and Matlab) are available from the corresponding authors upon request.

**Data availability:** The raw and treated data are available from the corresponding authors upon request.

## Actomyosin-driven force patterning controls endocytosis at the immune synapse Supplementary materials

### 1 Underestimation of the non-coordinated pool of forces

The non-coordinated displacements results of a force with a component perpendicular to the substrate. The traction force algorithm used in this work solves the 2D problem. In this paragraph we estimate the error we might make applying the TFM algorithm on a displacement that in reality results from a point-like force **F** = (0,0, *F*)*δ*(*x*)*δ*(*y*)*δ*(*z*). This force, perpendicular to the gel and exerted from a protrusion, (but the argument works also for outward forces, changing the sign), will generate a displacement **u** = **G_3_ · F** where **G**_3_ is the 3D Boussinesq Green function:

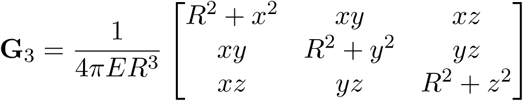

where *R*^2^ = *x*^2^ + *y*^2^ + *z*^2^. The displacement reads:

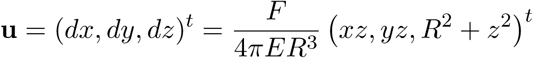

(where the *t* apex indicate transposed vectors). However, we measure only ũ = (*dx*, *dy*)^*t*^, hence the apparent components measured by TFM algorithm is 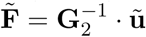 where

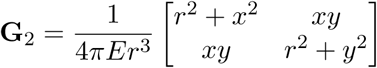

where *r*^2^ = *x*^2^+*y*^2^ indicate the position where we measure the displacement. Simplifying the equation we are left with

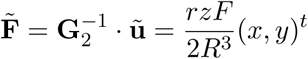

The ratio between force magnitude will give the percentage of underestimation:

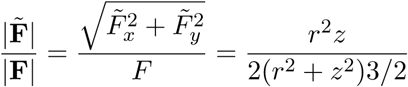

This function has a maximum in 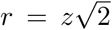 but is close to zero for small r, making our estimation completely wrong. The value of the maximum is independent of *z* and is 3^−3/2^ ≈ 20% meaning that at our best we underestimate the real force of a factor 5 (only when the displacement is measure close to the application point). For this reason we cannot rely on the TFM algorithm for 3D analysis. However the systematic underestimation is independent of the position and the quantification remains meaningful for spatial distribution and relative value *on average*.

### 2 Analysis of the coordinated and non-cordinated pool of forces

To distinguish between coordinated and non-coordinated pool we analyse each beads according to the degree of correlation with their neighbours. Each bead *i* is displaced of a vector 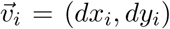. So for a chosen bead *j* we compute the correlation coefficient *r*_*j*_ with the *N* nearest neighbours belonging to region *C* as:

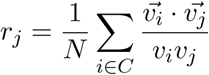

where *C* is the circle r radius 1 *μm* around bead *j*; this is the distance allowing to average on a mean of 5 beads. This index is equivalent to the average cosine of the angle formed by the displacements (in analogy to liquid crystal order parameter):

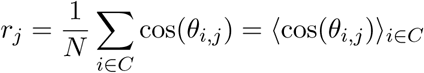

At each time point and for each beads *i* the value *r*_*i*_ is computed and the beads classified according to:

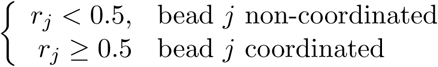

As can be deduced from the definition, *r*_*j*_ does not take into account the magnitude of displacement. Beads that, because of noisy detection and microscopy acquisition, show small apparent displacements will be included in this pool because the orientations of their displacement vectors are randomly distributed. This is the reason for the number of non-coordinated beads decreasing in Fig. 3d.

### 3 Myosin II driven pulsatile contractions leads to central patterns

In Reference [2] a model for cell motility is proposed where the velocity of the cell is coupled to the displacement of polarity cues via an actomyosin flow. Inspired by this model we propose that the pulsatile dynamics of actomyosin can transport membrane bound molecules towards the center of the synapse and and therefore help concentrating the endocytic machinery there. However the formation of a pattern, such as the accumulation of molecules in the center, is guaranteed only for certain range of parameters. Here we investigate whether in which condition actomyosin pulsatile contraction can generate patterns.

Let us consider a molecular species having concentration *c*(**r**, *t*) at the point **r** at instant *t*. We assume that, because of direct or indirect interactions with actin filaments, it is advected by the actin flow of speed **V**(*t*) and denote by *D* the diffusion coefficient. Its dynamics then follows

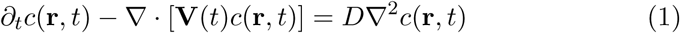

In our case **V** is time independent and we assume it is switched on/off with rate *k*_1_, *k*_0_ (according to a random telegraph process). For the sake of simplicity, we will assume below that the actin flow is centripetal and uniform in space, so that it can be written **V** = — *V*(*t*)**u**_*r*_ where **u**_*r*_ is the radial unit vector. Under these assumptions eq. 1 can be rewritten as a system of partial differential equations that describe the dynamics of the two populations *c*_1_ and *c*_0_ of particles subject or not to advection (respectively). The velocity is now time independent:

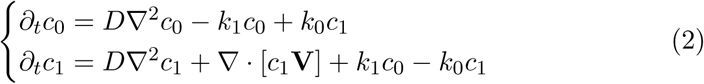

This system of linear differential equations can be solved explicitely for centripetal flows; it is however instructive to consider an adiabatic approximation where the typical timescales for diffusion time and advection are larger than the on-off kinetics of the velocity:

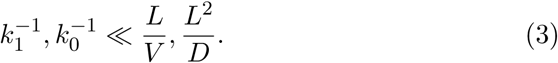

This is justified since typical on/off times are 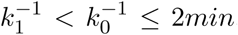, the size of the cell is *L* ≈ 5*μm* and diffusion is *D* ≈ 0.05*μm*^2^/*s*, which gives *L* ≈ 5*min* and *L*^2^/*D* ~ 10*min*.

In this case the concentrations equilibrate through the on/off kinetics faster than through the other processes and therefore one can assume *c*_0_/*c*_1_ = *k*_1_/*k*_0_ and substitute:

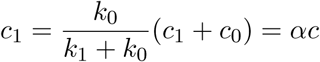

summing the two equations in (eq.2 and integrating, one obtains for the total concentration *c* = *c*_1_ + *c*_0_:

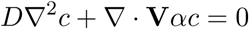

In cylindrical coordinates (and using the equation for the radial component (the system is centrosymmetric) reads:

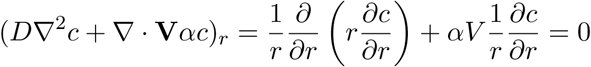

We solve this equation with open boundary conditions:

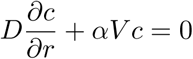

which gives the usual exponential solution:

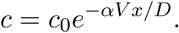

Clearly this defines a length scale *λ* = *D*/(*αV*) that can be used as a criterion for the formation of a pattern: when is smaller than the radius of the synapse *λ* ≪ *L* then the pattern is formed and there exists an appreciable gradient along the radius. Note that approximation in eq. 3 simplifies the problem but it is not necessary for its solution: being linear the stationary solution will be anyway exponential and the relevant length would be *λ* as above.

Pluging in experimental numbers:

- the diffusion constant of a molecule attached to the membrane is *D* ≤ 1*μm*^2^/*s* however for BCR this is typically *D* ≈ 0.05*μm*^2^/*s*
- flow speed is *V* ≈ 1*μm*/*s* from rough particle image velocimetry measurements on our data (in [1] a peak value 5 times higher is presented on stiff surfaces)
- *α* can be estimated observing that the typical time of a pulsation is few frames 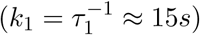 and the typical pulsation is around 170*s* 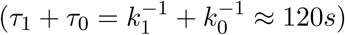: then 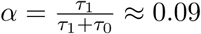

we find indeed a typical length scale in the problem of:

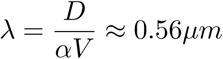

which means that the species are localised in the center provided that the diffusion is low enough (which is particularly true for membrane bound species, indeed *D* can be even smaller, for BCR this is typically *D*_*mem*_ = 0.05*μm*^2^/*s*). This means that the pattern will be destroyed on a timescale of *L*^2^/*D* ≈ 500*s*. Typical pulsations have shorter typical timescales, suggesting that this could be a mechanism to gather molecules in the center of the synapse.

### 4 Argument on the energy scale for affinity discrimination

We suggest also that the force patterning might play a role in improving affinity discrimination, in particular in reducing the noise in the center of the synapse. Why should be this important to isolate mechanically the region where the antigen is extracted?

In [3] the authors propose that the affinity discrimination is mechanical, this implies that the cell has to pull on the antigen to overcome an energetic barrier of height Δ*U*. The affinity depends essentially on the dissociation constant which scales exponentially with the energy:

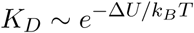

(where *k*_*B*_*T* 4_*p*_*N*.*nm*, the scale of thermal fluctuations at equilibrium).

A *N* folds increase in affinity between antigen 1 and 2 with two energetic barrier Δ*U*_1_ and Δ*U*_2_ can be written as:

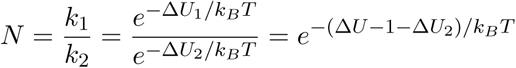

This results in an energy barrier difference of:

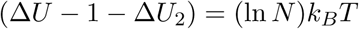

In the case of antigens used in [3] *N* = 10 and the energy difference is of the order of *ln*(10)*k*_*B*_*T* ≈ 2.3*k*_*B*_*T*. As a term of comparison the maximal work that a single Myosin II molecule can exert is 2.5*k*_*B*_*T*. Therefore the affinity discrimination mechanism could be easily disturbed by application of external forces or even weak mechanical noise. The peripheral stress could serve precisely to seal the synapse and mechanically isolate it from external noise.

### 5 Estimation of antigen extraction forces

In this section we give an estimation of the load on each HEL molecule based on our measurements. We first measured the concentration of HEL on gel using two techniques:

1. count of secondary antibodies fluorescence by using photobleaching and Poisson statistics as done in [4]; this gives about 12 fluorophores/*μm*^2^ which might means (assuming 1 fluorophore/secondary antibody, 2 primary antibodies/HEL and 2 secondary per primary) around maximum 50 HEL molecules/*μm*^2^.
2. measuring the amount of antigen not attached to the gel by Nanodrop and comparing with the original concentration. We found that when coating a gel with 100*g*/*ml* solution of HEL (7M), 30% of the proteins are retained, hence using 200l (i.e. ~ 20*μg* = 1.4*nMol* = 8.4 · 10^14^*molecules*) the gel retains 2.8 · 10^14^*molecules*. The mesh size of a 500*Pa* gels is about 50 – 100*nm* and this would allow the HEL molecule (~ 5*nm*) to go through the gel and functionalise it even in bulk. The surface of the gel is *A* = 250*mm*^2^ and its volume *V* = 25*mm*^3^ we obtain that there are roughly 11000*molecules/μm*^3^ or at the surface there are *c*^2/3^ = 500*molecules/μm*^2^.

We deduced that HEL concentraion is in the range 50 ~ 500*molecules/μm*^2^. We chose to consider the smallest value. The shear stresses we measure in our experiments are maximum 80*Pa*, hence a force of 100*pN/m*^2^. If all the forces were equally distributed among the HEL molecules one expects on average less than 2*pN* per molecule which make statistically difficult for shear force to contribute to antigen detachment. In contrast pointlike forces are necessary localised and reach values that could provide detachment of a single molecule. In fact mechanics is required for detachment forces over 12*pN* ([5]).

